# The ancestral endosymbiont *Blattabacterium* was lost ten times independently in Blattellidae, Pseudophyllodromiidae and Anaplectidae cockroaches

**DOI:** 10.64898/2026.08.23.746292

**Authors:** Zhuli Cheng, Yukihiro Kinjo, Esra Kaymak, David C.F. Rentz, Nathan Lo, Frédéric Legendre, Jan Šobotnik, Thomas Bourguignon

## Abstract

Most cockroaches and the termite *Mastotermes darwiniensis* are associated with *Blattabacterium*, an ancient obligate endosymbiont that participates in the nitrogen metabolism of its host. *Blattabacterium* has been vertically transmitted since it was acquired by the common ancestor of cockroaches and termites and was reportedly lost twice, once in the cockroach genus *Nocticola* and once in all termites except *Mastotermes darwiniensis*. Here, we acquired cockroach specimens spanning most of the cockroach phylogenetic tree to study *Blattabacterium* using shotgun sequencing. We found no traces of *Blattabacterium* in 64 specimens from ten independent lineages of cockroaches across three families: Blattellidae, Pseudophyllodromiidae, and Anaplectidae. The absence of *Blattabacterium* was confirmed with three PCR amplifications targeting the 16S and 23S ribosomal genes with primers specific to *Blattabacterium*. Notably, cockroaches lacking *Blattabacterium* were often infected by *Rickettsia* and *Wolbachia*, many of which were related to the mutualistic *Wolbachia* strain of *Cimex lectularius*, the common bed bug. These results indicate that cockroaches from Blattellidae, Pseudophyllodromiidae and Anaplectidae have lost their ancestral *Blattabacterium* endosymbiont at least ten times independently, with many of these losses possibly facilitated and compensated by new associations with mutualistic *Wolbachia* strains that may help provision the host with B vitamins.

**Significance statement:** Cockroaches harbor a beneficial bacterium with which they have been living together for hundreds of millions of years. The bacterium provides the host cockroaches with essential amino acids and vitamins. Here we discovered that this bacterium is absent in ten cockroach lineages. This absence might be compensated by the replacement of this bacterium with other bacteria that can provide essential nutrients, especially vitamins.

## Introduction

*Blattabacterium cuenoti* (hereafter: *Blattabacterium*) is the ancient obligate intracellular symbiont of cockroaches, inhabiting bacteriocytes, a specialized cell type in the cockroach fat body (Brooks 1970; Tokuda et al. 2008; López-Sánchez et al. 2009; Noda et al. 2020). *Blattabacterium* has been vertically transmitted since it was acquired by the common ancestor of termites and cockroaches and has since coevolved with its host (Bandi et al. 1995; Patiño-Navarrete 2013; Patiño-Navarrete et al. 2013; Arab et al. 2020; Kinjo et al. 2021). The endosymbiont is necessary for the proper development of its cockroach host, as its experimental removal reduces growth and fertility (Brooks and Richards 1955). Metabolic reconstruction of the *Blattabacterium* genome indicates that the endosymbiont is specialized in the recycling of nitrogenous wastes, converting urea into ammonia and providing its host with essential amino acids and vitamins (Sabree et al. 2009; Sabree et al. 2012; Kinjo et al. 2021; Kinjo et al. 2022).

*Blattabacterium* genomes are highly conserved, with all strains sharing a similar synteny with small deletions across the chromosome and very few inversions (Patiño- Navarrete et al. 2013; Kinjo et al. 2018; Kinjo et al. 2021). Despite this conserved synteny, *Blattabacterium* genomes slowly lose genes over millions of years, often in clusters representing entire metabolic pathways. For example, the *Blattabacterium* strains associated with the termite *Mastotermes darwiniensis* and with the wood-feeding cockroach *Cryptocercus*, the sister group of termites, have lost multiple pathways involved in the biosynthesis of essential amino acids (Tokuda et al. 2013; Kinjo et al. 2018). Similarly, the *Blattabacterium* strains associated with the soil-burrowing cockroaches *Geoscapheus dilatatus* and *Geoscapheus robustus* have lost genes involved in the biosynthesis of branched- chain amino acids and tryptophan (Beasley-Hall et al. 2024). Some *Blattabacterium* strains have experienced more intense genome reduction, especially those associated with species belonging to two cockroach families, Pseudophyllodromiidae and Anaplectidae, which harbor some of the smallest known *Blattabacterium* genomes, as small as 511 kbp, approximately 20% smaller than the 645 kbp of the largest known *Blattabacterium* genomes (Kinjo et al. 2021). These strains have lost up to 130 genes since they diverged from the strain infecting the last common ancestor of cockroaches (Kinjo et al. 2021). Gene loss in these strains is pervasive across COG functional categories; however, it is particularly pronounced in the COG categories for Coenzyme transport and metabolism (category H), Amino acid transport and metabolism (category E), and Cell wall/membrane/envelope biogenesis (category M) (Kinjo et al. 2021). These strains possess the most severely degraded *Blattabacterium* genomes described to date, a degradation characterized by the pervasive loss of metabolic functions associated with increased substitution rates and reduced GC content (Kinjo et al. 2021).

Despite being an obligate nutritional mutualist, *Blattabacterium* has been independently lost at least twice across Blattodea. The endosymbiont was lost once in the common ancestor of all termites except *Mastotermes darwiniensis*, the lineage sister to all other termites, possibly reflecting the acquisition of lignocellulolytic gut microbes capable of supplementing the termite diet through atmospheric nitrogen fixation and recycling of nitrogenous waste products (Sabree et al. 2012; Kinjo et al. 2018). The other documented independent loss of *Blattabacterium* occurred in *Nocticola*, a cockroach genus comprising many cave-dwelling species characterized by reduced eyes, reduced pigmentation, and accelerated rates of molecular evolution as compared to most other cockroaches (Roth 1988; Lo et al. 2007; Kovacs et al. 2024).

Here, we studied cockroach specimens using shotgun sequencing and identified 64 specimens belonging to ten independent cockroach lineages that, like non-*Mastotermes* termites and *Nocticola* spp., lack *Blattabacterium* contigs in the shotgun assemblies of their host fat bodies. We confirmed the absence of the endosymbiont with three PCR reactions carried out with *Blattabacterium*-specific primers. Two other endosymbionts, *Wolbachia* and *Rickettsia*, were also commonly detected across cockroaches, especially in cockroaches lacking *Blattabacterium*, suggesting endosymbiont replacement.

## Results

### Absence of Blattabacterium in 64 cockroach specimens

We analysed metagenome assemblies of cockroach specimens and found no *Blattabacterium contigs* in 64 specimens containing mitochondrial sequences that allowed us to position them on the cockroach phylogeny. We confirmed the presence or absence of *Blattabacterium* DNA in these 64 samples using PCR amplification and Sanger sequencing of three *Blattabacterium* rRNA loci. Gel electrophoresis revealed very faint bands for most of the 64 samples, contrasting with clear bands observed for the positive controls. These results suggest that most samples contain *Blattabacterium* DNA at extremely low concentrations, only revealed after 35 to 40 PCR cycles. We sequenced PCR products and found that the amplified sequences matched other samples that we processed in the same dissection and DNA extraction batches. Most of the amplicon sequences mapped to the sample processed immediately before the sample in question, indicating that the *Blattabacterium* sequences detected in the 64 samples lacking *Blattabacterium* contigs in the shotgun assemblies were cross-contaminations introduced during sample processing. We were able to trace the source of contamination for all but four samples, which all yielded amplicon sequences matching another cockroach family, indicating that these sequences also represented contaminations rather than genuine *Blattabacterium* presence. Details on PCR and Sanger sequencing results are available in Supplementary Data 1.

### Phylogenetic position of ten cockroach lineages lacking Blattabacterium

We reconstructed a phylogenetic tree using the first and second codon positions of 134 cockroach mitochondrial genomes, including 106 mitochondrial genomes sequenced in this study (Figure 1). The phylogenetic tree grouped the 64 samples lacking *Blattabacterium* into ten distinct lineages, with each lineage representing at least one independent loss of the endosymbiont. Among the ten lineages, four belonged to Anaplectidae, four to Pseudophyllodromiidae, and two to Blattellidae. In one lineage in Anaplectidae (lineage 3), we found that their conspecific close relative contains *Blattabacterium*, suggesting a recent loss of the endosymbiont.

**Figure 1.**
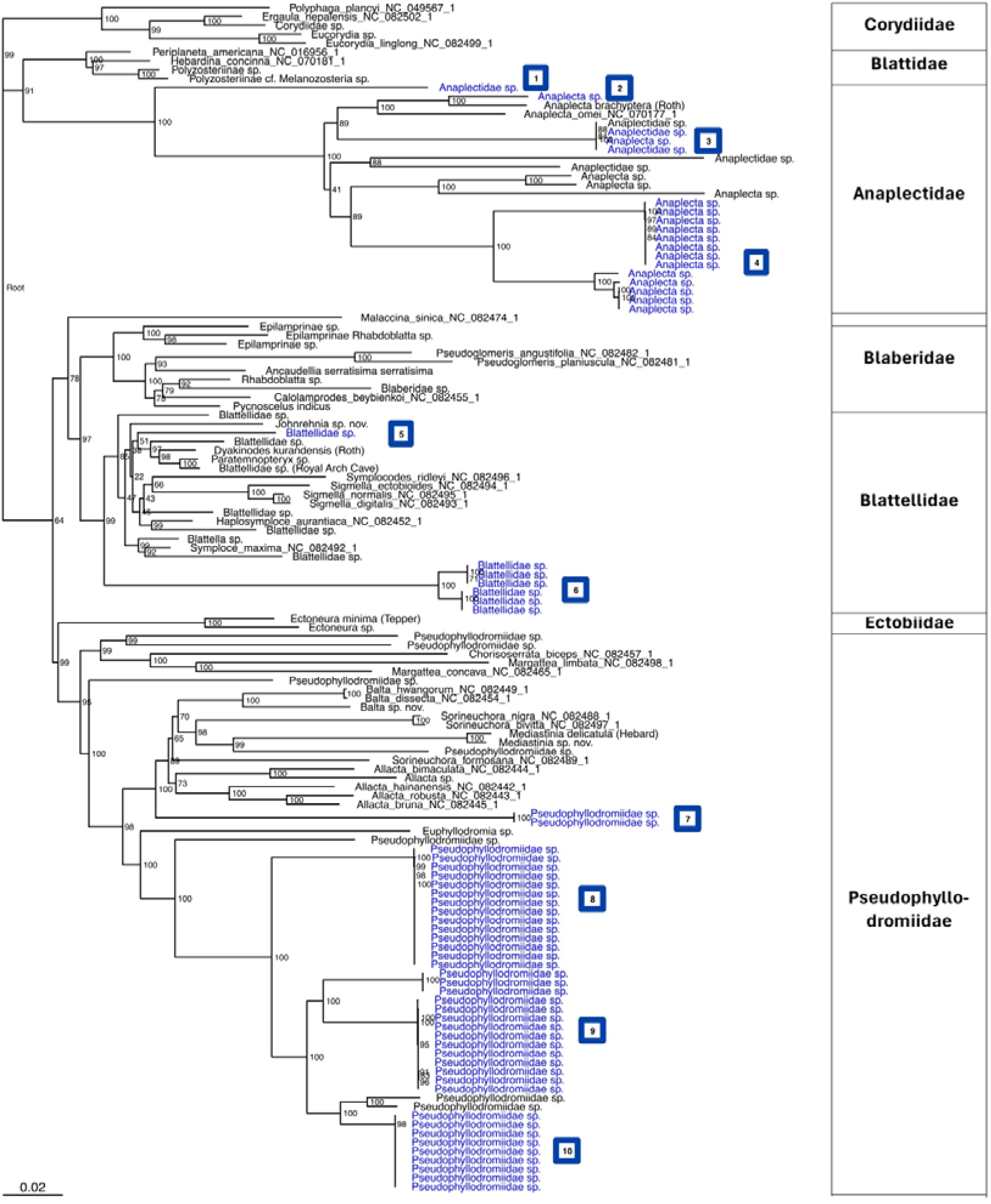
Maximum-likelihood phylogeny of cockroaches harboring or lacking *Blattabacterium*. The tree was reconstructed using the first and second codon positions of mitochondrial protein-coding genes. Cockroach specimens lacking *Blattabacterium* are indicated with blue tip labels. Lineages with independent loss of *Blattabacterium* are numbered from one to ten. Cockroach specimens harboring *Blattabacterium* are indicated with black tip labels. Sequences downloaded from NCBI RefSeq are indicated with NC numbers. Branch supports were assessed using 1,000 ultrafast bootstrap replicates. **Alt text:** This is a phylogenetic tree of 134 cockroach mitochondrial sequences from seven families, including Corydiidae, Blattidae, Anaplectidae, Blaberidae, Blattellidae, Ectobiidae and Pseudophyllodromiidae. 28 sequences are downloaded from NCBI RefSeq and 106 samples are generated in this study. Among the 106 samples, 64 samples do not contain *Blattabacterium*. These 64 samples are grouped into ten lineages with four in Anaplectidae, two in Blattellidae, and four in Pseudophyllodromiidae.

Among the ten lineages lacking *Blattabacterium*, lineage 7 (Pseudophyllodromiidae; two samples) was collected in Wanang Conservative Area in Papua New Guinea. Lineage 2 (Anaplectidae; one sample) was a dried specimen of National Museum of Natural History of France. The remaining eight lineages (61 samples) were collected in Ebogo, Cameroon.

### *Most cockroaches lacking* Blattabacterium *harbor* Wolbachia *and* Rickettsia

We searched for *Wolbachia* and *Rickettsia* across the 106 metagenome assemblies generated in this study. We detected contigs assigned to *Wolbachia* and *Rickettsia* in many metagenome assemblies, especially those from Pseudophyllodromiidae, Anaplectidae, and Blattellidae (Figure 2). *Wolbachia* reads were detected in 62 of the 106 samples, representing up to 7.07% of the metagenome reads. *Rickettsia* reads were detected in 56 samples, making up as much as 2.87% of the metagenome reads (Supplementary Data 2). Because dissection methods were variable among samples, consisting of fat body only in large specimens and whole abdomen in small specimens, our result likely underestimated the prevalence of *Wolbachia* and *Rickettsia*.

**Figure 2.**
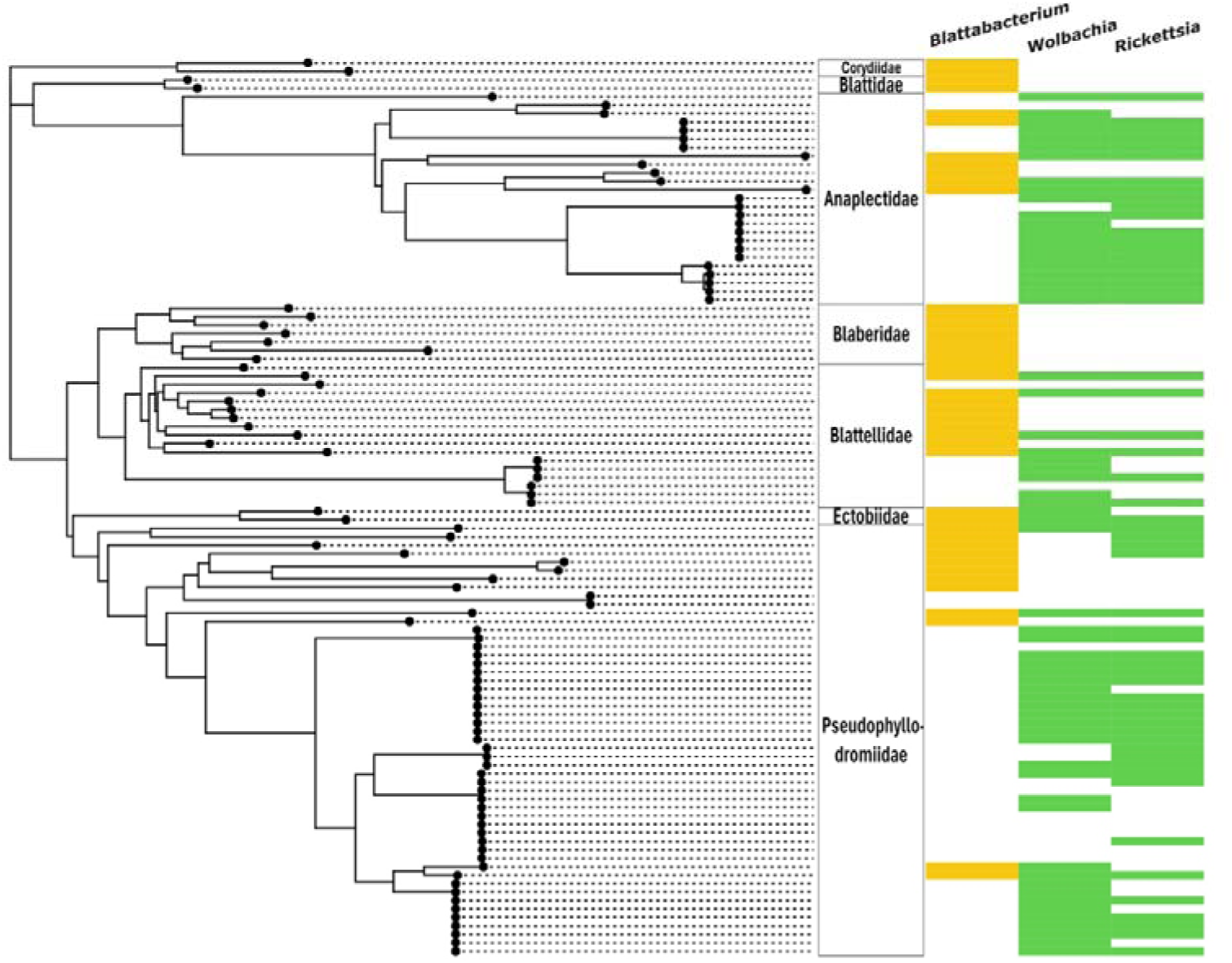
Presence and absence of *Blattabacterium*, *Wolbachia*, and *Rickettsia* mapped onto the cockroach host’s mitochondrial phylogeny. Colored cells indicate presence of the symbionts. **Alt text:** The figure shows a phylogeny of 106 samples generated in this study. Alongside the phylogeny, it is shown whether each sample contains *Blattabacterium*, *Wolbachia* and *Rickettsia*. Most samples that do not contain *Blattabacterium* contain *Wolbachia* and/or *Rickettsia*.

We identified *Wolbachia* and *Rickettsia* strains using BLASTn searches against the NCBI Nucleotide database. The closest reference strain to the *Wolbachia* contigs was that of the *Wolbachia* endosymbiont of *Cimex lectularius* (NZ_AP013028.1) in 40 of 62 samples containing *Wolbachia*. For the *Rickettsia* contigs, the closest reference strains were those of the *Rickettsia* endosymbionts of *Ceutorhynchus assimilis* (OU906081.1) and *Cimex lectularius* (CP084572.1) in 44 and 7 of 56 samples containing *Rickettsia*, respectively.

Notably, *Wolbachia* contigs from many samples of this study contained the genes for the biosynthesis of four B vitamins: biotin (*bioC*, *bioH*, *bioF*, *bioA*, *bioD*, *bioB*), folate (*folB*, *folK*, *folP*, *folC*, *folA*), riboflavin (*ribA*, *ribD*, *ribB*, *ribE*, *ribC*, *ribF*), and pyridoxine (*pdxJ*, *pdxH*). All known strains of *Blattabacterium* lack the genes involved in biotin biosynthesis and the genes involved in the biosynthesis of folate (B9), riboflavin (B2), and pyridoxine (B6) are frequently absent from the genomes of the *Blattabacterium* strains infecting Anaplectidae and Pseudophyllodromiidae (Figure S1, S2, S3).

## Discussion

Prior to this study, *Blattabacterium* loss was only reported in two lineages of Blattodea: the non-Mastotermitidae termites and the cockroach genus *Nocticola* (Lo et al. 2007; Sabree et al. 2012; Kinjo et al. 2018). Both groups are ecologically divergent from other cockroaches. Termites are eusocial and their ancestral diet consists of lignocellulose (Chouvenc et al. 2021), while most cockroaches are gregarious and opportunistic omnivores (Bell et al. 2007). *Nocticola* spp. are small cockroaches with pale coloration and variably reduced eyes and wings, comprising many cave-dwelling species that often feed on bat excrement (Roth 1988; Lo et al. 2007). In this study, we report the independent loss of *Blattabacterium* in ten additional cockroach lineages belonging to Pseudophyllodromiidae, Anaplectidae, and Blattellidae, expanding the known number of *Blattabacterium* loss across Blattodea.

Unlike is the case for termites and *Nocticola* spp., we know very little about the ecology of the cockroaches reported here to lack *Blattabacterium*, partly because Anaplectidae and Pseudophyllodromiidae are poorly studied cockroaches, and partly because most specimens used in this study were nymphs, preventing precise taxonomic identification. In contrast, we have genetic data informing us about the process of *Blattabacterium* loss. A first characteristic shared by the lineages lacking *Blattabacterium* is their comparatively long branches on the mitochondrial phylogeny, indicating elevated substitution rates. Anaplectidae and Pseudophyllodromiidae have accelerated substitution rates in comparison to other cockroach families sampled in this study. Blattellidae do not show higher substitution rates in comparison to other cockroaches, but lineage 6 lacking *Blattabacterium* has substantially longer branches than their close relatives. This pattern mirrors the elevated substitution rates previously documented in *Nocticola* spp., especially for the mitochondrial protein-coding genes (Legendre et al. 2015; Kovacs et al. 2024), suggesting that elevated substitution rates in the host cockroach mitochondrial genome are linked to *Blattabacterium* loss.

A second characteristic is the affiliation of most cockroach lineages lacking *Blattabacterium* with Anaplectidae and Pseudophyllodromiidae, two cockroach families harboring the *Blattabacterium* strains with the most reduced genomes. The loss of *Blattabacterium* therefore primarily occurred in cockroaches hosting *Blattabacterium* that already experienced significant degradation by gene loss, a pattern that often precedes the extinction of obligate endosymbionts (McCutcheon et al. 2019).

A third factor potentially associated with *Blattabacterium* loss is the acquisition of new replacing symbionts. The presence of *Wolbachia* has previously been reported in multiple cockroach families, including Ectobiidae, Pseudophyllodromiidae, Blattellidae, and Blaberidae (Lo, 2002; Vaishampayan et al. 2007; Choubdar et al. 2023; Oladipupo et al. 2023; Guse and Pietri 2024). In our study, we detected contigs assigned to *Wolbachia* or *Rickettsia* in numerous samples, particularly in Anaplectidae, Blattellidae, and Pseudophyllodromiidae. Notably, the *Wolbachia* strain most closely related to the majority of our contigs is the strain of *Cimex lectularius*, an obligate nutritional mutualist vertically transmitted from mother to offspring and whose main functions include the biosynthesis of B vitamins, especially biotin and riboflavin (Hosokawa et al. 2010; Nikoh et al. 2014; Moriyama et al. 2015). All *Blattabacterium* strains studied to date lack the genes of the biotin operon and therefore are unable to synthesize biotin (Kinjo et al. 2021; Kinjo et al. 2022). In addition, *Blattabacterium* strains associated with Pseudophyllodromiidae and Anaplectidae often lost the ability for de novo biosynthesis of folate (B9), riboflavin (B2), and pyridoxine (B6), three vitamins that the *Blattabacterium* strains of other cockroaches can synthesize (Kinjo et al. 2021). These results are in line with the negative correlation between *Blattabacterium* and *Wolbachia* abundance in *Supella longipalpa* (Guse and Pietri 2024), suggesting that the loss of *Blattabacterium* is associated with the recruitment of *Wolbachia* as a mutualistic endosymbiont. This is reminiscent of the recruitment of parasites as mutualists in other symbiotic systems, such as the recurrent recruitment of *Ophiocordyceps* fungi as a replacement of *Hodgkinia* endosymbionts in cicadas (Matsuura et al. 2018). Whether the acquisition of *Wolbachia* facilitated relaxed selection and the loss of *Blattabacterium* or *Wolbachia* were recruited as mutualistic endosymbionts following the loss of *Blattabacterium* is unclear and require additional experiments.

Because Blattellidae generally host *Blattabacterium* that did not experience significant genome reduction since they were vertically inherited from the last common cockroach ancestor, the loss of *Blattabacterium* by two lineages of Blattellidae is unexpected. It is possible that shifts in nitrogen metabolism contributed to *Blattabacterium* loss in some Blattellidae lineages. Cockroaches are unusual among insects for storing uric acid instead of excreting it, enabling nitrogen recycling via *Blattabacterium*. However, three species in Blattellidae, *Shawella couloniana*, *Symploce hospes*, and *Parcoblatta* spp., have been reported to void uric acid in formed pellets (Cochran 1973; Mullins and Cochran 1976). Moreover, *Shawella couloniana* and *Symploce hospes* store relatively little uric acid internally even when fed a protein-rich diet, suggesting limited nitrogen recycling (Mullins and Cochran 1976). These cockroaches may not rely heavily on uric acid storage and recycling, limiting their dependence on *Blattabacterium*, which could lead to relaxed selection in *Blattabacterium*.

Finally, another characteristic that the specimens lacking *Blattabacterium* share is that almost all of them (61 out of 64) were collected in Ebogo in Cameroon. We are not sure whether the geographical location has biased our detection of *Blattabacterium* loss and whether the geographical location is linked to relaxed selection in the endosymbiont, such as by relaxing requirements on nitrogen recycling or via increased opportunity for symbiont replacements.

## Conclusion

We identified ten lineages of cockroaches belonging to Pseudophyllodromiidae, Anaplectidae, and Blattellidae that have independently lost *Blattabacterium*, substantially increasing the number of lineages of Blattodea known to have lost *Blattabacterium*. These results improve our understanding of the coevolutionary history between cockroaches and their *Blattabacterium* endosymbionts, showing that elevated substitution rates in cockroach mitochondrial genomes and prior gene loss in *Blattabacterium* preceded the loss of the endosymbiont. In addition, the recruitment of mutualistic *Wolbachia* strains may have compensated for the loss of *Blattabacterium* in these cockroach lineages. Overall, our results show that cockroaches frequently lose their *Blattabacterium*, as is the case in other insect endosymbionts (McCutcheon et al. 2019).

## Material and Methods

### Specimens

This project used 106 cockroach specimens from seven families: Blattellidae, Blaberidae, Pseudophyllodromiidae, Ectobiidae, Anaplectidae, Blattidae, and Corydiidae. Most specimens were collected directly from the field in Cameroon (83 specimens), Australia (8 specimens), Japan (3 specimens), Papua New Guinea (8 specimens), and China (1 specimens) and stored in ethanol or RNAlater at −20 °C until DNA extraction. We also used 3 dried pinned specimens from the National Museum of Natural History in Paris, France. These specimens were stored at room temperature until DNA extraction. Specimens were identified based on morphological characters whenever possible; however, many specimens could not be identified morphologically, especially nymphs, and were identified based on mitochondrial sequences. Collection localities and species identifications are provided in Supplementary Data 3.

### Shotgun sequencing

Fat bodies were dissected using forceps and scalpels under a ZEISS Stemi 508 stereo microscope (Carl Zeiss, Germany). For small specimens from which insufficient fat body tissues could be obtained, we used the whole abdomen. Genomic DNA was extracted from dissected tissues using the DNeasy Blood & Tissue Kit (Qiagen, Hilden, Germany). Library preparation was performed using the NEBNext Ultra II FS DNA Library Prep Kit for Illumina (New England Biolabs, Ipswich, MA, USA) following a modified protocol. For samples preserved in ethanol or RNAlater, we followed the manufacturer’s protocol, except for the reagent volumes, which were decreased to one-fifteenth of the recommended volumes to reduce costs. For dried museum specimens, in addition to using one-fifteenth of the reagent volumes, we omitted the enzymatic fragmentation step for the library preparation. Other library preparation steps were performed as instructed by the manufacturer’s protocol. Libraries were sequenced in-house on a NovaSeq 6000 system (Illumina, San Diego, CA, USA) generating 2 × 151 bp paired-end reads.

### Genome assembly

We used FaQCs v2.10 (Lo and Chain 2014) to filter and trim raw reads using the parameters “-q 25 -min_L 120 -avg_q 30 -adaptor 1”. Metagenome assembly was performed using SPAdes v3.13.0 (Nurk et al. 2017; Prjibelski et al. 2020) and TCSF-IMRA v2.7.2 (Kinjo et al. 2015). The assembly quality was subsequently improved using Pilon v1.23 (Walker et al. 2014).

### *Presence/absence of* Blattabacterium, Wolbachia*, and* Rickettsia

We identified contigs of three symbionts, *Blattabacterium*, *Wolbachia*, and *Rickettsia*, using BLASTn searches against the NCBI nucleotide database using ncbi-blast v2.13.0 (Camacho et al. 2009). A symbiont was considered absent from a metagenome assembly when we were unable to identify any contig derived from that symbiont. We estimated the relative abundance of each identified symbiont by mapping reads to assembled contigs using Bowtie2 (Langmead and Salzberg 2012) to estimate the proportion of reads assigned to each symbiont. Read depth was estimated using SAMtools (Li et al. 2009).

### Phylogenetic reconstruction of cockroaches

We built a cockroach phylogenetic tree using the protein-coding genes of mitochondrial genomes. Cockroach mitochondrial sequences were retrieved from our metagenome assemblies using BLASTn searches against the NCBI nucleotide database using ncbi-blast v2.13.0. Our dataset comprised 106 samples from this study, including 64 samples lacking *Blattabacterium* and 42 samples selected from our broader collection of samples from which we assembled complete circular *Blattabacterium* chromosomes. We supplemented this dataset with 28 complete mitochondrial genomes of cockroaches downloaded from NCBI RefSeq to place our dataset in a broader phylogenetic context. We annotated mitochondrial sequences with MitoFinder v1.4 (Jühling et al. 2011; Allio et al. 2020) using 1333 blattodean mitochondrial sequences from the NCBI Nucleotide database as references. We extracted the nucleotide sequences of all 13 protein-coding genes from the annotations, discarding genes fragmented across multiple contigs and pseudogenes that contained more than one stop codon. We used MAFFT v7.505 (Katoh and Standley 2013) with the options “--globalpair -- maxiterate 1000” to align the amino acid sequences of each gene independently. Amino acid alignments were back-translated into nucleotide alignments using pal2nal v14.1 (Suyama et al. 2006). Alignment positions with more than 2% gaps were removed from the nucleotide alignments with trimAl v1.4.rev22 (Capella-Gutiérrez et al. 2009). The trimmed nucleotide alignments were concatenated, and the third codon positions were removed to minimize substitution saturation using FASconCAT-G v1.04 (Kück and Longo 2014). We converted the four-nucleotide alignments into binary alignments of pyrimidine and purine to avoid potential issues caused by GC content heterogeneity (Ishikawa 2012). Maximum-likelihood phylogenetic trees were reconstructed with IQ-TREE v2.0.7(Chernomor et al. 2016; Hoang et al. 2018; Wang et al. 2018; Minh et al. 2020), with the first and second codon positions treated as separate partitions. The best-fit substitution model was selected for each partition using ModelFinder. A GTR2+FO+I+G4 model was selected for the first codon position, and a GTR2+FO+R5 model for the second codon position. Branch support was assessed using 1,000 ultrafast bootstrap replicates (Minh et al. 2013).

### Validation of Blattabacterium absence with PCR and Sanger sequencing

Our shotgun sequencing approach flagged 64 samples as potentially lacking *Blattabacterium* and containing a sufficient number of mitochondrial contigs to place the specimen on the phylogenetic tree of cockroaches. We verified the absence of *Blattabacterium* using three PCR reactions with three pairs of primers specific to *Blattabacterium*. We designed two pairs of primers targeting the *Blattabacterium* 16S rRNA gene and one pair of primers targeting the *Blattabacterium* 23S rRNA gene. We also used a pair of primers targeting the cockroach 18S rRNA gene as a positive control. To maximize the PCR amplification of *Blattabacterium* markers, we used a touchdown PCR program with a wide range of annealing temperatures (Table S1).

Our primers were designed based on a broader dataset of metagenome assemblies of 446 cockroach specimens. More specifically, we identified the ten cockroach specimens that were the closest relatives of the 64 target samples based on mitochondrial phylogeny and used these specimens as positive controls. We verified that all four pairs of primers aligned to the ten references with a maximum of one mismatch located in the central region of the primer sequences to maximize binding efficiency. Primer specificity to *Blattabacterium* was confirmed by aligning primer sequences against 45 Flavobacteriia sequences downloaded from NCBI, ensuring that multiple mismatches were present across each primer sequence. Pilot experiments confirmed that the three pairs of *Blattabacterium* primers and one pair of cockroach primers yielded amplicons in the ten reference samples.

PCR amplifications were performed on the 64 samples for which no *Blattabacterium* contigs were assembled from our shotgun sequencing data using a Mastercycler® X50s (Eppendorf, Hamburg, Germany). Each PCR reaction was performed in a volume of 20 µL containing 0.5 µL of each primer (10 µM), 10 µL of 2× QuickTaq HS Dye Mix (Toyobo, Osaka, Japan; DTM-101), and 2-4 µL of DNA extract. The total volume was adjusted to 20 µL with Milli-Q water. PCR conditions consisted of an initial denaturation step at 94 °C for 2 min, followed by 35 or 40 cycles of denaturation at 94 °C for 30 sec, touchdown annealing for 30 sec (Table S1), and extension at 68 °C for 1.5 min, with a final extension at 72 °C for 2 min. Negative controls, consisting of Milli-Q water instead of DNA extracts, were also included on each PCR plate. For each PCR amplification, 3 µL of PCR product was visualized on 1.5% agarose gels (NIPPON GENE, Tokyo, Japan) prepared in TAE buffer and run for about 50 minutes at 100V on an i-MyRun II system (Cosmo Bio Co., Ltd., Tokyo, Japan). The lengths of the bands were estimated by comparison with 3-6 µL of DNA ladder spanning 100-1500 bp (Toyobo, Osaka, Japan; DNA-035). We imaged the gels using an iBright FL1500 Imaging System (Invitrogen, Waltham, MA, USA).

PCR products were purified using AMPure XP magnetic beads at a 1.8x ratio (Beckman Coulter, Brea, CA, USA; A63881) and eluted in 20 µL of UltraPure™ DNase/RNase-Free Distilled Water (Invitrogen, Waltham, MA, USA; 10977015). Purified PCR products were quantified using the Qubit™ 1× dsDNA High Sensitivity Assay Kit (Thermo Fisher Scientific, Waltham, MA, USA; Q33231). Sanger sequencing was performed using the BigDye™ Terminator v3.1 Cycle Sequencing Kit (Applied Biosystems, Tokyo, Japan; 4337454). Each sequencing reaction was performed in a total volume of 10 µL containing 1.5 µL of 5× sequencing buffer, 0.4 µL of BigDye reaction mix, 2-5 µL (100ng) of purified PCR product, and 0.5 µL of each primer (0.5 µM concentration). The volume was adjusted to 10 µL with nuclease-free water. Sequencing was carried out under the following conditions: an initial denaturation at 96°C for 1 min, followed by 25 cycles of denaturation at 96°C for 10 sec, annealing at 50°C for 5 sec, and extension at 60°C for 4 min. Sequencing products were analyzed by capillary electrophoresis on a SeqStudio Genetic Analyzer (Life Technologies Japan Ltd., Tokyo, Japan). Forward and reverse sequencing reactions were performed independently. The obtained sequences were trimmed at both ends using the trimfq function of seqtk v1.5 (Li 2025) with default settings. Forward and reverse trimmed sequences were queried against *Blattabacterium* genomes sequenced in the same batch using BLASTn to trace the identity of amplified sequences.

## Acknowledgements

We thank members of the Environmental Science and Information Section at OIST, Sizuma Yanagisawa, and Zuzana Varadínová Kotyk for sharing specimens. We thank the OIST’s Sequencing Section for the sequencing services and the OIST’s Scientific Computing and Data Analysis Section for providing access to the OIST computing cluster. We also thank the Bioinformatics User Group for installing and maintaining Bioinformatics tools on the OIST’s High Performance Computing System. We thank members of the Evolutionary Genomics Unit at OIST for helpful discussions. We acknowledge financial support from Okinawa Institute of Science and Technology Graduate University, with subsidy funding from the Cabinet Office, Government of Japan,

## Use of AI

ChatGPT (OpenAI) was used to assist in writing and debugging R/Python/bash scripts for data processing and visualization. The author verified the correctness of all code and analytical outputs.

## Data availability

All sequences generated in this study will be made available on GenBank. Supplementary Data 1, 2, and 3 can be downloaded at https://doi.org/10.6084/m9.figshare.32515245.

## Author Contributions

Z.C., Y.K., and T.B. conceptualized the experiments. D.C.F.R., N.L., F.L., and J.S. obtained the cockroach specimens. Z.C., Y.K., and E.K., performed lab experiments and generated the raw data. Z.C. and Y.K. performed the bioinformatic analyses. Z.C. and T.B. wrote the original draft. All authors edited and accepted the final version of this manuscript.

## References

Allio R et al. 2020. MitoFinder: Efficient automated large-scale extraction of mitogenomic data in target enrichment phylogenomics. Mol Ecol Resour. 20(4):892–905 [accessed 2026 Apr 21]. /doi/pdf/10.1111/1755-0998.13160. 10.1111/1755-0998.13160;JOURNAL:JOURNAL:14718286;WGROUP:STRING:PUBLICATION

Arab DA et al. 2020. Evolutionary rates are correlated between cockroach symbionts and mitochondrial genomes. Biol Lett. 16(1) [accessed 2022 Jun 20]. /pmc/articles/PMC7013487/. 10.1098/rsbl.2019.0702

Bandi C et al. 1995. The Establishment of intracellular symbiosis in an Ancestor of Cockroaches and Termites. Proc Biol Sci. 259(1356):293–299 [accessed 2022 Oct 3]. 10.1098/rspb.1995.0043

Beasley-Hall PG et al. 2024. Shrinking in the dark: Parallel endosymbiont genome erosions are associated with repeated host transitions to an underground life. Insect Sci. 31(6):1810 [accessed 2026 Mar 2]. https://pmc.ncbi.nlm.nih.gov/articles/PMC11632294/. 10.1111/1744-7917.13339

Bell WJ, Roth LM, Nalepa CA. 2007. Cockroaches, Ecology, Behavior and Natural History. The Johns Hopkins University Press.

Brooks MA. 1970. Comments on the classification of intracellular symbiotes of cockroaches and a description of the species. J Invertebr Pathol. 16:249–258 [accessed 2023 Jul 20]

Brooks MA, Richards AG. 1955. Intracellular symbiosis in cockroaches. I. Production of aposymbiotic cockroaches. Biol Bull. 109(1):22–39 [accessed 2026 Mar 13]. /doi/pdf/10.2307/1538656?download=true. 10.2307/1538656

Camacho C et al. 2009. BLAST+: architecture and applications. BMC Bioinformatics 2009 10:1. 10(1):421- [accessed 2026 Apr 21]. https://link.springer.com/article/10.1186/1471-2105-10-421. 10.1186/1471-2105-10-421

Capella-Gutiérrez S, Silla-Martínez JM, Gabaldón T. 2009. trimAl: a tool for automated alignment trimming in large-scale phylogenetic analyses. Bioinformatics. 25(15):1972 [accessed 2023 Oct 21]. /pmc/articles/PMC2712344/. 10.1093/BIOINFORMATICS/BTP348

Chernomor O, Von Haeseler A, Minh BQ. 2016. Terrace Aware Data Structure for Phylogenomic Inference from Supermatrices. Syst Biol. 65(6):997–1008 [accessed 2026 Apr 21]. 10.1093/sysbio/syw037. 10.1093/SYSBIO/SYW037

Choubdar N et al. 2023. Wolbachia infection in native populations of Blattella germanica and Periplaneta americana. PLoS One. 18(4):e0284704 [accessed 2026 Mar 26]. https://pmc.ncbi.nlm.nih.gov/articles/PMC10118093/. 10.1371/journal.pone.0284704

Chouvenc T, Šobotník J, Engel MS, Bourguignon T. 2021. Termite evolution: mutualistic associations, key innovations, and the rise of Termitidae. Cell Mol Life Sci. 78(6):2749 [accessed 2026 May 27]. https://pmc.ncbi.nlm.nih.gov/articles/PMC11071720/. 10.1007/S00018-020-03728-Z

Cochran DG. 1973. Comparative analysis of excreta from twenty cockroach species. Comp Biochem Physiol A Physiol. 46(2):409–419 [accessed 2026 Mar 18]. 10.1093/aesa/46.4.547. 10.1016/0300-9629(73)90429-5

Guse K, Pietri JE. 2024. Endosymbiont and gut bacterial communities of the brown-banded cockroach, Supella longipalpa. PeerJ. 12:e17095 [accessed 2026 Mar 26]. https://pmc.ncbi.nlm.nih.gov/articles/PMC10959106/. 10.7717/PEERJ.17095

Hoang DT et al. 2018. UFBoot2: Improving the Ultrafast Bootstrap Approximation. Mol Biol Evol. 35(2):518–522 [accessed 2026 Apr 21]. 10.1093/molbev/msx281. 10.1093/MOLBEV/MSX281

Hosokawa T et al. 2010. Wolbachia as a bacteriocyte-associated nutritional mutualist. Proc Natl Acad Sci U S A. 107(2):769–774 [accessed 2026 Apr 14]. /doi/pdf/10.1073/pnas.0911476107?download=true. 10.1073/PNAS.0911476107;WEBSITE:WEBSITE:PNAS-SITE;WGROUP:STRING:PUBLICATION

Jühling F et al. 2011. Improved systematic tRNA gene annotation allows new insights into the evolution of mitochondrial tRNA structures and into the mechanisms of mitochondrial genome rearrangements. Nucleic Acids Res. 40(7):2833 [accessed 2026 Apr 21]. https://pmc.ncbi.nlm.nih.gov/articles/PMC3326299/. 10.1093/NAR/GKR1131

Katoh K, Standley DM. 2013. MAFFT multiple sequence alignment software version 7: improvements in performance and usability. Mol Biol Evol. 30(4):772–780 [accessed 2022 Apr 20]. https://academic.oup.com/mbe/article/30/4/772/1073398. 10.1093/MOLBEV/MST010

Kinjo Y et al. 2018. Parallel and gradual genome erosion in the Blattabacterium endosymbionts of Mastotermes darwiniensis and Cryptocercus wood roaches. Genome Biol Evol. 10(6):1622–1630. 10.1093/gbe/evy110

Kinjo Y et al. 2021. Enhanced mutation rate, relaxed selection, and the “domino effect” are associated with gene loss in Blattabacterium, A cockroach endosymbiont. Mol Biol Evol. 38(9):3820–3831. 10.1093/molbev/msab159

Kinjo Y et al. 2022. Coevolution of metabolic pathways in Blattodea and their Blattabacterium endosymbionts, and comparisons with other insect-bacteria symbioses. Microbiol Spectr. 10(5) [accessed 2023 Oct 18]. https://journals.asm.org/doi/10.1128/spectrum.02779-22. 10.1128/SPECTRUM.02779-22/SUPPL_FILE/SPECTRUM.02779-22-S0002.XLS

Kinjo Y, Saitoh S, Tokuda G. 2015. An efficient strategy developed for next-generation sequencing of endosymbiont genomes performed using crude DNA isolated from host tissues: A case study of Blattabacterium cuenoti inhabiting the fat bodies of cockroaches. Microbes Environ. 30(3):208–220. 10.1264/jsme2.ME14153

Kovacs TGL et al. 2024. Dating in the Dark: Elevated Substitution Rates in Cave Cockroaches (Blattodea: Nocticolidae) Have Negative Impacts on Molecular Date Estimates. Syst Biol. 73(3):532–545 [accessed 2026 Mar 30]. 10.1093/sysbio/syae002. 10.1093/sysbio/syae002

Kück P, Longo GC. 2014. FASconCAT-G: extensive functions for multiple sequence alignment preparations concerning phylogenetic studies. Front Zool. 11(1):81 [accessed 2026 Apr 21]. https://pmc.ncbi.nlm.nih.gov/articles/PMC4243772/. 10.1186/S12983-014-0081-X

Langmead B, Salzberg SL. 2012. Fast gapped-read alignment with Bowtie 2. Nature Methods 2012 9:4. 9(4):357–359 [accessed 2026 Apr 23]. https://www.nature.com/articles/nmeth.1923. 10.1038/nmeth.1923

Legendre F et al. 2015. Phylogeny of Dictyoptera: Dating the origin of cockroaches, praying mantises and termites with molecular data and controlled fossil evidence. PLoS One. 10(7):e0130127 [accessed 2023 Oct 20]. https://journals.plos.org/plosone/article?id=10.1371/journal.pone.0130127. 10.1371/JOURNAL.PONE.0130127

Li H et al. 2009. The Sequence Alignment/Map format and SAMtools. Bioinformatics. 25(16):2078–2079 [accessed 2026 Apr 23]. 10.1093/bioinformatics/btp352. 10.1093/BIOINFORMATICS/BTP352

Li H. 2025. Seqtk: Toolkit for processing sequences in FASTA/Q formats.

Lo C, Chain PSG. 2014. Rapid evaluation and quality control of next generation sequencing data with FaQCs. BMC Bioinformatics. 15(1):1–8 [accessed 2023 Oct 21]. https://bmcbioinformatics.biomedcentral.com/articles/10.1186/s12859-014-0366-2. 10.1186/S12859-014-0366-2/TABLES/3

Lo N et al. 2007. Cockroaches that lack Blattabacterium endosymbionts: the phylogenetically divergent genus Nocticola. Biol Lett. 3(3):327 [accessed 2022 Oct 5]. /pmc/articles/PMC2464682/. 10.1098/RSBL.2006.0614

López-Sánchez MJ et al. 2009. Evolutionary Convergence and Nitrogen Metabolism in Blattabacterium strain Bge, Primary Endosymbiont of the Cockroach Blattella germanica. PLoS Genet. 5(11):1000721 [accessed 2023 Feb 10]. http://genome.jgi-psf. 10.1371/journal.pgen.1000721

Matsuura Y et al. 2018. Recurrent symbiont recruitment from fungal parasites in cicadas. Proc Natl Acad Sci U S A. 115(26):E5970–E5979 [accessed 2023 Mar 31]. https://www.pnas.org/doi/abs/10.1073/pnas.1803245115. 10.1073/PNAS.1803245115/SUPPL_FILE/PNAS.1803245115.SAPP.PDF

McCutcheon JP, Boyd BM, Dale C. 2019. The Life of an Insect Endosymbiont from the Cradle to the Grave. Current Biology. 29(11):R485–R495 [accessed 2023 Mar 6]. 10.1016/J.CUB.2019.03.032

Minh BQ et al. 2020. IQ-TREE 2: New Models and Efficient Methods for Phylogenetic Inference in the Genomic Era. Mol Biol Evol. 37(5):1530–1534 [accessed 2026 Apr 21]. 10.1093/molbev/msaa015.10.1093/MOLBEV/MSAA015

Minh BQ, Nguyen MAT, Von Haeseler A. 2013. Ultrafast Approximation for Phylogenetic Bootstrap. Mol Biol Evol. 30(5):1188–1195 [accessed 2022 Sep 6]. https://academic.oup.com/mbe/article/30/5/1188/997508. 10.1093/MOLBEV/MST024

Moriyama M, Nikoh N, Hosokawa T, Fukatsu T. 2015. Riboflavin provisioning underlies wolbachia’s fitness contribution to its insect host. mBio. 6(6) [accessed 2026 Apr 14]. /doi/pdf/10.1128/mbio.01732-15?download=true. 10.1128/MBIO.01732-15;JOURNAL:JOURNAL:MBIO;CTYPE:STRING:JOURNAL

Mullins DE, Cochran DG. 1976. A comparative study of nitrogen excretion in twenty-three cockroach species. Comp Biochem Physiol A Physiol. 53(4):393–399. 10.1016/S0300-9629(76)80162-4

Nikoh N et al. 2014. Evolutionary origin of insect-Wolbachia nutritional mutualism. Proc Natl Acad Sci U S A. 111(28):10257–10262 [accessed 2026 Apr 14]. /doi/pdf/10.1073/pnas.1409284111?download=true. 10.1073/PNAS.1409284111;ISSUE:ISSUE:DOI

Noda T et al. 2020. Bacteriocytes and Blattabacterium Endosymbionts of the German Cockroach Blattella germanica, the Forest Cockroach Blattella nipponica, and Other Cockroach Species. 102108/zs200054. 37(5):399–410 [accessed 2022 Jul 15]. https://bioone.org/journals/zoological-science/volume-37/issue-5/zs200054/Bacteriocytes-and-Blattabacterium-Endosymbionts-of-the-German-Cockroach-Blattella-germanica/10.2108/zs200054.full. 10.2108/ZS200054

Nurk S, Meleshko D, Korobeynikov A, Pevzner PA. 2017. MetaSPAdes: A new versatile metagenomic assembler. Genome Res. 27(5):824–834 [accessed 2023 Oct 21]. https://genome.cshlp.org/content/27/5/824.full. 10.1101/GR.213959.116/-/DC1

Oladipupo SO et al. 2023. The prevalence of Wolbachia in multiple cockroach species and its implication for urban insect management. J Econ Entomol. 116(4):1307–1316 [accessed 2026 Mar 26]. 10.1093/jee/toad098. 10.1093/jee/toad098

Patiño-Navarrete R. 2013. Evolutionary genomics and functional studies on the metabolic role of Blattabacterium, primary endosymbiont of cockroaches.

Patiño-Navarrete R, Moya A, Latorre A, Peretó J. 2013. Comparative genomics of Blattabacterium cuenoti: The frozen legacy of an ancient endosymbiont genome. Genome Biol Evol. 5(2):351–361 [accessed 2023 Jul 20]. 10.1093/gbe/evt011. 10.1093/GBE/EVT011

Prjibelski A et al. 2020. Using SPAdes De Novo Assembler. Curr Protoc Bioinformatics. 70(1):e102 [accessed 2022 Aug 30]. https://onlinelibrary.wiley.com/doi/full/10.1002/cpbi.102. 10.1002/CPBI.102

Roth LM. 1988. Some cavernicolous and epigean cockroaches with six new species, and a discussion of the Nocticolidae (Dictyoptera- Blattaria). Revue suisse de zoologie. (95):297– 321. 10.5962/bhl.part.79654

Sabree ZL et al. 2012. Genome shrinkage and loss of nutrient-providing potential in the obligate symbiont of the primitive termite Mastotermes darwiniensis. Appl Environ Microbiol. 78(1):204–210 [accessed 2023 Sep 8]. https://journals.asm.org/journal/aem. 10.1128/AEM.06540-11

Sabree ZL, Kambhampati S, Moran NA. 2009. Nitrogen recycling and nutritional provisioning by Blattabacterium, the cockroach endosymbiont. Proc Natl Acad Sci U S A. 106(46):19521–19526 [accessed 2023 Feb 14]. https://www.pnas.org/doi/abs/10.1073/pnas.0907504106. 10.1073/PNAS.0907504106/SUPPL_FILE/0907504106SI.PDF

Suyama M, Torrents D, Bork P. 2006. PAL2NAL: robust conversion of protein sequence alignments into the corresponding codon alignments. Nucleic Acids Res. 34(suppl_2):W609– W612 [accessed 2023 Oct 21]. 10.1093/nar/gkl315. 10.1093/NAR/GKL315

Tokuda G et al. 2008. Purification and partial genome characterization of the bacterial endosymbiont Blattabacterium cuenoti from the fat bodies of cockroaches. BMC Res Notes. 1(1):1–9 [accessed 2022 Oct 3]. https://bmcresnotes.biomedcentral.com/articles/10.1186/1756-0500-1-118. 10.1186/1756-0500-1-118/TABLES/1

Tokuda G et al. 2013. Maintenance of essential amino acid synthesis pathways in the Blattabacterium cuenoti symbiont of a wood-feeding cockroach. Biol Lett. 9(3) [accessed 2026 Mar 2]. 10.1098/rsbl.2012.1153

Vaishampayan PA et al. 2007. Molecular evidence and phylogenetic affiliations of Wolbachia in cockroaches. Mol Phylogenet Evol. 44(3):1346–1351 [accessed 2026 Mar 26]. 10.1371/journal.ppat.0020043. 10.1016/j.ympev.2007.01.003

Walker BJ et al. 2014. Pilon: An integrated tool for comprehensive microbial variant detection and genome assembly improvement. PLoS One. 9(11):e112963 [accessed 2023 Oct 21]. https://journals.plos.org/plosone/article?id=10.1371/journal.pone.0112963. 10.1371/JOURNAL.PONE.0112963

Wang HC, Minh BQ, Susko E, Roger AJ. 2018. Modeling Site Heterogeneity with Posterior Mean Site Frequency Profiles Accelerates Accurate Phylogenomic Estimation. Syst Biol. 67(2):216–235 [accessed 2026 Apr 21]. 10.1093/sysbio/syx068. 10.1093/SYSBIO/SYX068

